# Cannabis use is associated with sexually dimorphic changes in executive control of visuospatial decision-making

**DOI:** 10.1101/2020.01.08.899526

**Authors:** Parker J. Banks, Patrick J. Bennett, Allison B. Sekuler, Aaron J. Gruber

## Abstract

When the outcome of a choice is less favourable than expected, humans and animals typically shift to an alternate choice option. Several lines of evidence indicate that this “lose-shift” responding is an innate sensorimotor response strategy that is normally suppressed by executive function. Therefore, the lose-shift response provides a covert gauge of cognitive control over choice mechanisms. We report here that the spatial position, rather than visual features, of choice targets drives the lose-shift effect. Furthermore, the ability to inhibit lose-shift responding to gain reward is different among male and female habitual cannabis users. Increased self-reported cannabis use was concordant with suppressed response flexibility and an increased tendency to lose-shift in women, which reduced performance in a choice task in which random responding is the optimal strategy. On the other hand, increased cannabis use in men was concordant with reduced reliance on spatial cues during decision-making, and had no impact on the number of correct responses. These data (63,600 trials from 106 participants) provide strong evidence that spatial-motor processing is an important component of economic decision-making, and that its governance by executive systems is different in men and women who use cannabis frequently.

## Introduction

Adaptive decision-making is governed by the interaction of several brain circuits, each of which has unique aspects that are advantageous under particular circumstances. For instance, a classic distinction has been made between goal-directed control, involving the prefrontal cortex and medial striatum, and habitual control systems comprised of the sensorimotor cortex and lateral striatum (Balleine and O’Doherty, 2010; Gruber and McDonald, 2012). The goal-directed system appears to implement executive functions, such as working memory and strategic planning (Fuster, 1989; Passingham and Wise, 2012). However, cannabis abuse compromises the ability of this executive system to suppress sensorimotor responding (Knight et al, 1999; Rae et al, 2015; Malone and Taylor, 2006; Filbey and Yezhuvath, 2013). Here we report a new dissociation among executive and sensorimotor systems governing choice, which allows us to quantify their interaction while accounting for important confounding factors such as decision time and learning.

When rewards are uncertain, the most pervasive strategy in animals and humans is to repeat choices that have previously led to reward (win-stay), and to shift away from choices following reward omission (lose-shift) (Kamil and Hunter, 1970; Worthy et al, 2013; Thorndike, 2017). Although the win-stay and lose-shift are complementary behaviours, they are anatomically disassociated among goal-directed and sensorimotor systems. Lesions to the rodent lateral striatum (LS), which is homologous to the human putamen and essential for sensorimotor control (Parent and Hazrati, 1995), disrupt lose-shift responding but not winstay (Skelin et al, 2014; Gruber et al, 2017; Thapa and Gruber, 2018). A similar shifting deficit has been observed in humans with damage to putamen or insula (Danckert et al, 2011). Conversely, lesions of the rodent ventromedial striatum (VS), a key structure in goal-directed control that recieves inputs from prefrontal cortex (Voorn et al, 2004), disrupts win-stay but not lose-shift responding (Gruber et al, 2017). Several other behavioural features in rodents and humans support this anatomical disassociation. The win-stay and lose-shift exhibit different temporal dynamics (Gruber and Thapa, 2016), developmental trajectories (Ivan et al, 2018), and responses to reward feedback (Banks et al, 2018). Moreover, lose-shift responding (but not win-stay) drastically increases in adult humans under cognitive load, as well as in young children (Ivan et al, 2018). These data suggest that executive function can over-ride lose-shift responding, which can be characterized as a reflexive response by the sensorimotor striatum. This is consistent with a long history of research indicating that executive function can suppress reflexive or habitual motor responses (Chamberlain and Sahakian, 2007).

The LS/putamen receives prominent inputs from both the somatosensory and motor cortices (Brasted et al, 1999), and encodes the motor aspects of decision-making (Burton et al, 2015). Consequently, decisions and their associated motor actions are represented in egocentric (body-centred) spatial coordinates (Kesner and DiMattia, 1987; Palencia and Ragozzino, 2005). The dorsolateral caudate, which receives inputs from the dorsolateral PFC, is also necessary for egocentric spatial processing (Possin et al, 2017). The VS/nucleus accumbens encodes the value of choices and can engage a wide range of spatial-motor actions when executing a single decision involving abstract representations (Burton et al, 2015; Mashhoori et al, 2018). It encodes responses in both egocentric and allocentric (world-centred) spatial coordinates, likely involving its prominent inputs from the hippocampus and prefrontal cortex (PFC) (De Leonibus et al, 2005; Voorn et al, 2004; Possin et al, 2017). These data suggest that the control of actions by sensorimotor systems will be restricted to an egocentric framework heavily dependent on the position of targets, whereas the control by executive systems have the capacity to use abstract features of targets. This is supported by the dissociated effects of cannabis on neural structures and performance on spatial versus non-spatial tasks, as described next.

The recreational use of psychoactive substances has complex short-term and long-term effects on the brain, some of which dissociate. *Δ*^9^-tetrahydrocannabinol (THC) administration increases dopamine release in the LS, while the VS remains unaffected (Sakurai-Yamashita et al, 1989). Behaviourally, dopamine signalling in the dorsolateral striatum is necessary for normal spatial memory, motor control, and visospatial learning, while reward processing and goal-directed learning rely on dopamine signalling in the medial striatum (Darvas and Palmiter, 2009, 2010). This provides an explanation for the observation that THC reduces spatial processing and visuospatial memory in humans (Cha et al, 2007), particularly in females (Pope et al, 1997). Recreational drugs also differentially influence win-stay and lose-shift responding. THC and amphetamine cause large changes in lose-shift behaviour in rats and humans, while the win stay is only weakly affected (Wong et al, 2017b,a; Paulus et al, 2002).

Because the LS is necessary for lose-shift responding, and it processes information in egocentric coordinates, we hypothesized that lose-shift responses are calculated according to the position of a target relative to the participant, rather than other visual features of target identity. We further hypothesized that frequent cannabis use will disrupt the normal positional-dependence of lose-shift and the ability of executive systems to govern sensorimotor control. We tested these hypotheses while human participants were engaged in a competitive decision-making task between two choices. Crucially, the choices were visually distinct and changed their spatial configuration unpredictably between each trial. We found that following a loss, lose-shift behaviour was robustly associated with a choice’s previous location, rather than its visual identity. The win-stay was only weakly associated with previous choice position, and this association was eliminated by global changes in target position. Although female cannabis users exhibited reduced task performance and increased lose-shift responding, their reliance on spatial information was not different than controls. Male cannabis users, however, did exhibit a reduced reliance on spatial information. These data support the dissociation of choice among systems with different spatial propensities, and reveal a sexual dimorphism of recreational cannabis use and the function of these systems.

## Methods

### Behavioural Task

During the experiment, participants played a competitive game colloquially called “Matching Pennies” against a computer opponent. The task display consisted of two distinct targets (a blue circle and yellow square) presented on a 15″ touchscreen monitor (Figure 1). On each trial, participants would choose either target by touching it on the screen. They would then receive visual feedback indicating “You Win” or “You Lose” for 1.5s. On each trial, the computer algorithm attempted to predict which target would be selected. If the participant selected this target, the trial was a loss. Otherwise it was a win. The algorithm attempted to minimize the number of wins for participants. The optimal strategy for the participants is to be unpredictable in choice, in which case they win on 50% of trials. Because the winstay and lose-shift are predictable, subjects should learn to suppress these responses as the session progresses. This task provides measures of lose-shift, win-stay, and cognitive flexibility as participants adapted their choices to the computer opponent.

**Fig. 1.**
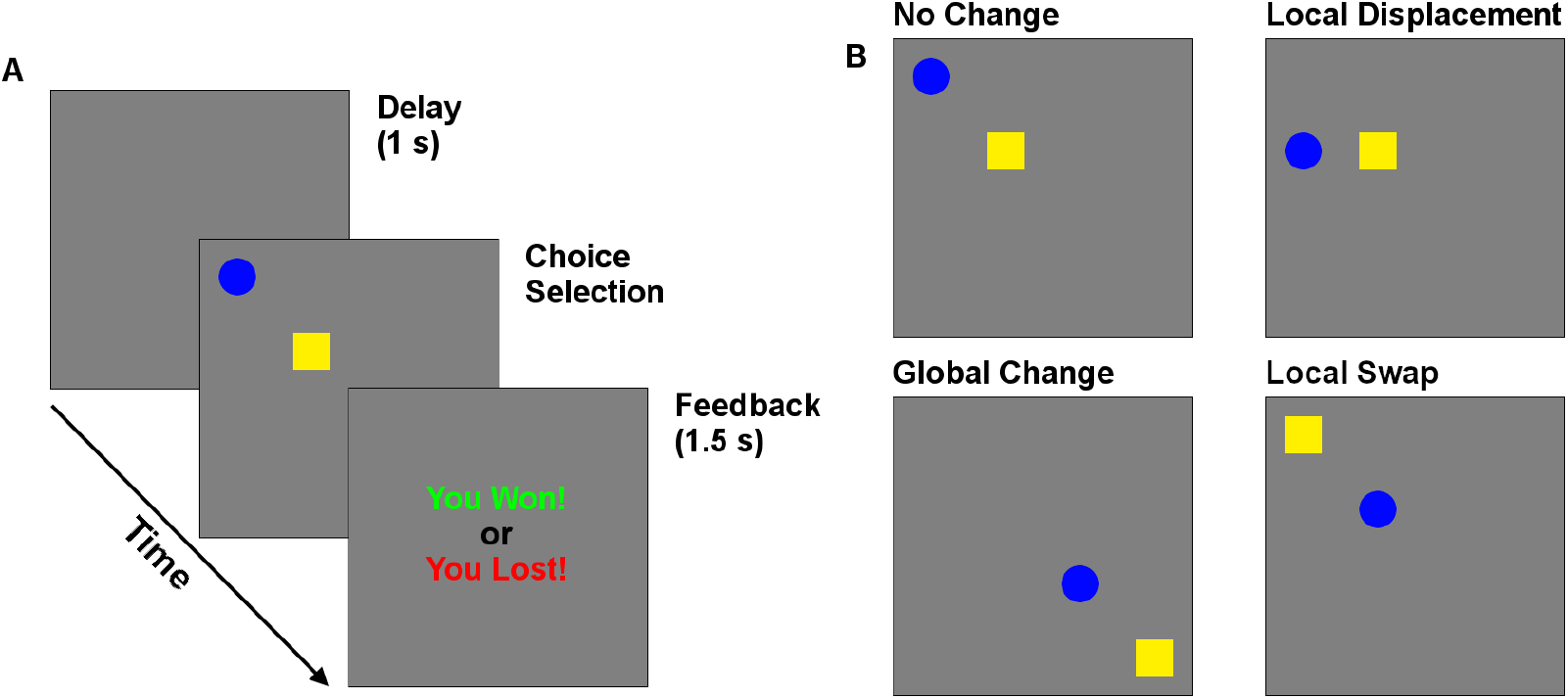
Behavioural task. (A) Timeline of trials in the matching pennies game. (B) Reconfiguration of targets, which could undergo local (swap & displace) and global changes in position.

The computer used four types of algorithms to detect patterns in (*i*) participants’ choices, (*ii*) switching from one choice to another, (*iii*) choices paired with rewards (e.g., blue square after a loss), & (*iv*) switching paired with rewards (e.g., swapping choices after a loss). Specifically, on each trial the computer examined a subject’s recent choice & reward history (e.g., shifting from the blue target to the yellow after a loss). The choice that most accurately predicted the subject’s past choice history was selected as the prediction of the present choice. Patterns of choices 1-6 trials in length were considered, resulting in 24 total prediction strategies. On each trial, the best performing strategy (computed over all previous trials in the session) was used to predict participants’ choices. If all strategies failed to beat the participant on ≥ 50% of past trials, the computer would select choices randomly.

The effect of cue position was investigated by moving the location of one or both cues from one trial to the next. The changes came in two types - local and global. The screen was divided into four equal quadrants, each of which contained an invisible 2×2 grid in its center. Local changes occurred within each grid, while global changes involved shifting among quadrants. Three local manipulations are of particular interest to investigate the importance of position and cue identity. The ‘Control’ case is when the cue positions remain in the same locations. The ‘Swap’ case occurs when the targets swap positions (a local change). The ‘Displace’ condition occurs when the previously selected choice moves to a previously empty position in the same 2×2 choice grid, while the other target remains in its previous position. Global changes occurred independently of each of these 3 local changes, for a total of 6 possible changes of target positions across subsequent trials, which were were selected randomly on each trial. This manipulation allowed us to determine how the position of targets relative to each other, and to the participants, affected choice following wins and losses. In particular, this design allows us to test if participants avoid the screen position of a target following a loss, as expected by the egocentric processing framework of LS. The alternative is they instead avoid the target regardless of position.

### Procedure

All procedures and experimental tasks were approved by the McMaster University Research Ethics Board. One hundred six undergraduates (53 males, mean age = 19.40, *SD* = 2.74) from McMaster University participated in the study in exchange for payment. After providing informed consent, participants played 600 trials of the task. They were informed that they would win nothing each time the computer predicted their choice and 3¢ each time it could not, rounded up to the nearest $5 upon completion of the experiment. Participants were given no guidance as to optimal decision-making strategies.

After task completion, participants were screened with the South Oaks Gambling Screen, the alcohol, smoking and substance involvement screening test (ASSIST) v3.0, Adult ADHD Self-Report Scale v1.1, and an additional demographic questionnaire. Habitual cannabis users were defined as those meeting the criteria for brief or intensive treatment (score > 3) on the ASSIST cannabis subtest. Total drug use was also recorded as the ASSIST score summed across all drug subtypes. Males had a mean ASSIST score of 19.11, with 32.08, 39.62, and 60.38% meeting the criteria for alcohol, cannabis, or any recreational drug use requiring intervention. Females had a mean ASSIST of 16.66, with 28.30, 30.19, and 45.28% meeting alcohol, cannabis, or general drug use criteria (See Table 1).

**Table 1.** Demographic data and ASSIST questionnaire scores (± SEM) for the studied groups.

| Sex | Group | N | Age | Cannabis | ASSIST |
| --- | --- | --- | --- | --- | --- |
| Female | Control | 37 | 19.2±2.0 | 0.3±.9 | 4.1±6.2 |
|  | Cannabis | 16 | 18.9±1.1 | 12.8±8.6 | 40.0±25.4 |
| Male | Control | 32 | 19.3±2.6 | 0.7±1.2 | 8.6±8.4 |
|  | Cannabis | 21 | 20.29±4.3 | 12.8±8.0 | 35.1±21.3 |

### Analysis

Participants’ responses were analyzed for proportion of lose-shift & win-stay responses, averaged over 5 blocks of 120 trials each, and conditioned on the type of cue shift relative to the previous trial’s target positions. As a measure of behavioural flexibility, the binary response entropy (*H*) for each participant was calculated from 4-trial choice sequences as:

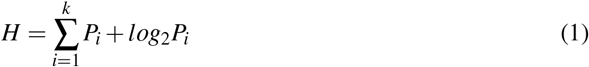

Where *P_i_* is the probability of each choice sequence, and *k* is the total number of sequences possible (i.e., 16). For example, a participant that exhibited the choice pattern “circle-square-circle-square” to the exclusion of all other patterns, would have an entropy of 0 bits, while a participant responding randomly would have an entropy of 4 bits. Response entropy and task performance were averaged over the experimental session for each participant. Decision times were measured as the time to make a response following presentation of the choice selection screen. They were normalized using the inverse transform (1/RT) and averaged after removing 131 erroneous RTs of <3 ms. The inverse transform was used to normalize RTs because it produced more normalized (Gaussian) distributions than did the log or square-root transforms.

The differences of marginal means of derived quantities (decision time, lose-shift, etc.) were tested by analysis of variance (for categorical factors) or co-variance (for continuous factors) using repeated-measure, mixed-effects models. Each model utilized a maximal random-effects structure and was fit in R using the lme4 package (Bates et al, 2014). A maximal model ensured that variations in effects between participants, and between trial blocks within each participant, were properly controlled (Barr et al, 2013). Degrees of freedom and *p*-values were calculated using the Welch-Satterthwaite equation and type-III sums of squares. The effects of local changes in position were assessed via planned paired t-tests comparing the effects of spatial swaps & displacement relative to the no-change condition.

We also used the *Q-learning with forgetting* (Barraclough et al, 2004) reinforcement learning model to examine the effects of cannabis use, local changes, and global changes on reward sensitivity, choice stochasticity, and learning rates. In this model the probability of selecting one of the two choices (*C*) on a given trial (*t*) are calculated according to the softmax equation (Sutton and Barto, 2018).
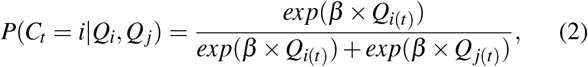

where *Q_i_* and *Q_j_* are the value each subject assigns to choices *i* and *j*. *β* refers to the inverse temperature that balances exploiting known action-reward associations with exploring more of the state/action space. As such, larger values of *β* indicate a greater tendency to choose the most highly valued action. The values of each choice are updated from rewards (*R*) according to the following rules:

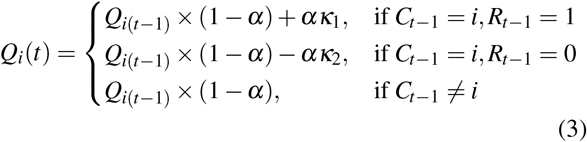

where *α* is the learning and forgetting rates for the chosen and unchosen action, *κ*_1_ is the strength of reinforcement from reward, and *κ*_2_ is the strength of aversion from failing to receive a reward. These three parameters were treated as stochastic variables that follow a random walk process. As such, they were free to vary throughout the experiment. Conversely, *β* was treated as a deterministic variable that remained fixed throughout the experiment. These parameters were fit for each subject using the VBA toolbox (Daunizeau et al, 2014).

To determine how local swaps, displacement, and global changes influenced RL parameters (i.e., hidden state values), we performed a Volterra decomposition of *α*, *κ*_1_, & *κ*_2_ values for each trial onto five basis functions (*u*): previous choice, outcome, local displacement, swap, and global change (relative to no change), according to Eq. 4:

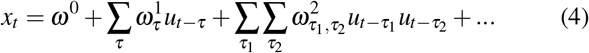

Volterra modelling allows for observation of input response characteristics of non-linear systems as Volterra weights (Boyd et al, 1984). At each trial *t* the Volterra weight *x* of a given parameter is estimated from inputs *u* over trials *t* to a lag of *τ* (set to 32 trials) using a series of Volterra kernels *ω*. The first kernel *ω*^1^ represents the linear transformation of lagged input basis functions into the output, *ω*^2^ represents the effect of past inputs being dependent on other earlier inputs, and so on. These weights provide a measure of how subjects’ valuation of each choice changes from baseline in response to past choices and outcomes. The benefit of Volterra modelling over analysis of raw prediction error is that the effect of current and past inputs on hidden state responses can be estimated. Inputs were also orthogonalized so that the effect of one input (e.g., local swaps) is computed independently of all other inputs (e.g., global changes). To control for trial order effects, we also detrended inputs prior to decomposition using a cubic polynomial.

## Results

### Recreational cannabis use coincides with sensorimotor dominance of decisions in women

Each of the 106 included participants performed 600 trials of the task, for a total of 63,600 trials in the dataset. We first sought to reveal how recreational cannabis use and biological sex affected overall performance on the task. We compared the effects of sex (male, female) and habitual cannabis use on task performance via a 2×2 analysis of variance (ANOVA) with type-III sums of squares and a zero-sum constraint. Cannabis use was associated with decreased task performance in females [*t*(51) = −3.123, *p* = .003, *d* = -.934]^*A*^, while males were unaffected (*p* = .777) (Fig. 2.A). Consequently, there was a significant main effect of cannabis use [*F*_1,102_ = 4.772, *p* = .032] and a sex × cannabis use interaction [*F*_1,102_ = 6.540, *p* = .012], while the main effect of sex was not significant (*p*=.271). This interaction between sex and cannabis use was reflected in similar trends in response entropy and decision times (Figs. 2.B & C). While the effects of sex, cannabis use, and sex × cannabis use on response entropy fell just short of significance (*p* >.055 in all cases), female cannabis users did exhibit significantly lower response entropy relative to controls [*t*(51)= −2.201, *p* = .032, *d* = -.658]^*B*^. Cannabis use was also associated with decreased decision times in females [*t*(51) = −3.024, *p* = .004, *d* = -.905]^*C*^, while those of men were again unaffected (*p*=.399), as indicated by a significant sex × cannabis use interaction [*F*_1,102_ = 7.701, *p* = .007]. However, no main effects were present (*p* > .112 in both cases).

**Fig. 2.**
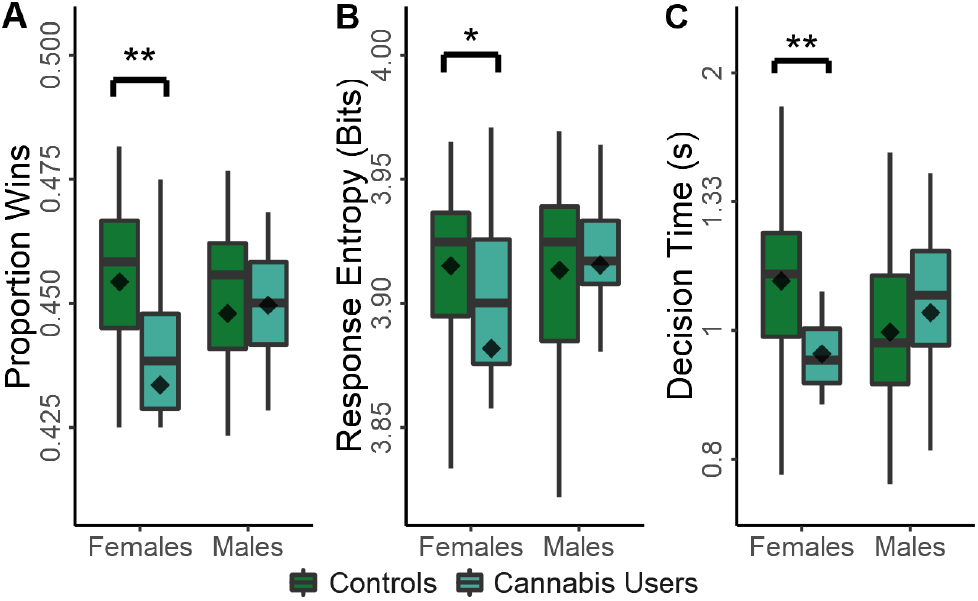
Effect of recreational drug use use on measures of task performance in males and females: proportion of wins (A); response entropy (B); and decision times (C). Plots show the conventional descriptive statistics: mean (diamond), median (horizontal line in the box), 25th/75th percentiles (box edges), and outliers (dots). Note: \**p* < .05, * * *p* < .01,

As expected, task performance was highly correlated with response entropy [*r*(104) = .699, *p* < .001], highly anti-correlated with mean lose-shift tendencies [*r*(104) = -.605, *p* < .001], and not correlated with win-stay responding [*r*(104) = -.095, *p* = .333]. Therefore, frequent cannabis use in females strongly coincides with increased lose-shift responding and decreased response times. These features are consistent with dominance of sensorimotor control in decision processes. Moreover, the tendency for increasingly stereotyped response sequences in females that frequently used cannabis further suggests a reduction in cognitive flexibility, defined here as a loss of ability to generate varied response types in response to many losses so as to improve task performance.

### Spatial cues drive lose-shift and win-stay responding

The optimal strategy on the task is to simply select targets at random on each trial of the session. Deviation from this optimal strategy is revealing of neural processes guiding behavioural choice. Lose-shift responding is maladaptive in this context, but nonetheless is a prevalent strategy. We next investigated to what extent spatial and/or other visual features of targets affect the propensity for lose-shift and win-stay behaviours. The task design allows us to test if participants avoid the screen position of a target following a loss, as expected by the egocentric processing framework of LS, or if they avoid the target itself regardless of position. We compared the effects of local & global changes in target position via 2 (global change, no change) × 3 (no local change, displace, swap) mixed-effects models with repeated measures and a full random effect structure (see Methods). We then conducted planned comparisons of marginal means for significant effects revealed by ANOVA.

Lose-shift behaviour was strongly affected by local changes in position across the 600 trials of the session parsed into 5 blocks [RMANOVA, *F*_2,117.43_ = 25.643, *p* < .001]. No effects of global changes [*F*_1,320.75_ = .368, *p* = .545], nor a local × global interaction [*F*_2,147.87_ = 2.285, *p* = .105] were present. As seen in the left panel of Figure 3.A, participants exhibited a high degree of lose-shift responding when choices did not move between trials, particularly in the first 3 blocks (360 trials). However, swapping choice positions strongly reversed their associated lose-shift probabilities [*t*(529) = −7.249, *p* < .001, *d* = -.507]^*D*^. This is particularly evident in the first 3 blocks. Because we are computing shifts with respect to each target (rather than position), a lose-shift probability less than 0.5 on swap trials indicates that participants are selecting the same target, but in a new location. This is a lose-stay response in terms of target identity, but a lose-shift in terms of spatial position. In other words, the blue and orange lines should overlap if lose-shift is computed with respect to target identity irrespective of location. Therefore, lose-shift is based on the previous position of an unrewarded target, rather than its identity as distinguished by other features (color and shape).

**Fig. 3.**
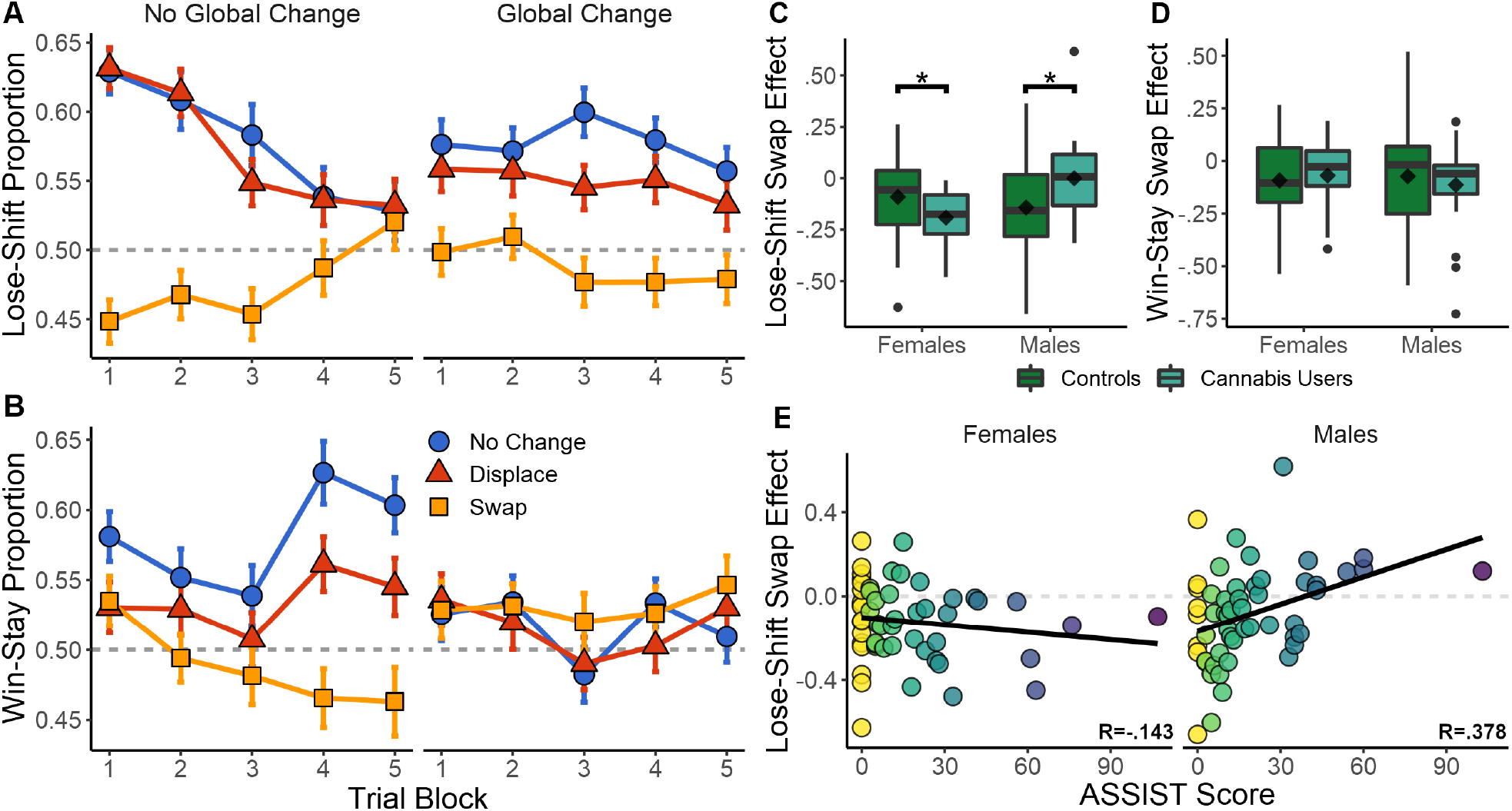
Effects of target reconfiguration and recreational drug use on reinforcement-driven behaviour. A-B: Effect of local & global changes in choice position on lose-shift and win-stay tendencies for all participants. SEM in error bars. C-D: Box plots of the difference in lose-shift and win stay when target positions are swapped as compared to no change. The effect of swapping targets on lose-shift is higher in women who use cannabis, but lower in men who use cannabis, than their sex-matched controls. E: Correlation between total drug use (ASSIST) and swap effect on lose-shift.

Although participants are able to eventually suppress lose-shift responding after hundreds of trials, this only occurs in the absence of global changes in target positions (compare left and right panels of 3.A). The effects of local swaps remained even in the presence of global changes [*t*(529) = −8.056, *p* < .001, *d* = -.459]^*E*^. Furthermore, the effect of local displacement was only significant in the presence of global changes [*t*(529) = −2.827, *p* = .005, *d* = -.160]^*F*^, and reduced the participant’s use of the lose-shift as compared to global changes and no displacement. Global changes thus appear to immediately reduce the probability of lose-shift early in the session, but to also interfere with the ability of participants to eventually learn to suppress this sub-optimal response near the end of the session.

The same analysis was repeated for win-stay responses. As seen in Figure 3.B, local changes [*F*_2,105.65_ = 5.470, *p* = .005] and a local × global interaction [*F*_2,159.33_ = 11.070, *p* < .001] had a significant effect on win-stay responding, while the main effect of global changes fell short of significance [*F*_1,1117.7_ = 3.792, *p* = .052]. Across the entire session, both local swaps [*t*(529) = −6.565, *p* < .001, *d* = -.438]^*G*^ and displacement [*t*(529) = −4.388, *p* < .001, *d* = -.221]^*H*^ reduced win-stay responding when no global change was present. Unlike the lose-shift, win-stay behaviour was initially unaffected by local changes. As trial blocks progressed, however, both win-stay and the effects of local changes increased. Global changes completely eliminated any effects of local changes throughout the session. The most parsimonious explanation is that subjects eventually learn to suppress the shift response, which reveals the stay response as the session progresses. To test if these are in competition, we computed the correlation of lose-shift and win-stay using data separated into the 5 trial blocks in the no change condition. Initially they were uncorrelated [*r*(104) = .008, *p* = .935]. However, as trials progressed the win-stay and lose-shift exhibited an increasingly negative correlation of [*r*(104) = -.310, *p* = .001] in block 3 and [*r*(104) = -.406, *p* < .001] in block 5. Therefore, competition between these strategies increases with time such that the competition is initially biased toward shift responses, but becomes biased toward stay responses as the session progresses.

Because the time between reinforcement and subsequent decisions affect lose-shifting (Gruber and Thapa, 2016; Ivan et al, 2018), we next analyzed whether changes in these response types could be explained by effects of target manipulation on decision times. Analysis of decision times (inverse transformed) indicated significant effects of local [*F*_2,140.14_ = 9.668, *p* < .001] and global [*F*_1, 109.81_ = 29.731, *p* < .001] changes in position, and a local × global interaction [*F*_2,140.15_ = 9.672, *p* < .001]. To further elucidate these effects, Table 2 provides decision times for each response type following local and global changes. Regardless of response type, global changes significantly increased decision times. More importantly, local swaps increased the time of lose-shift, win-shift, & win-stay responses, particularly when no global changes were present. While it could be argued that this increase is only due to the extra time needed to move to a new location, the fact that these effects are not consistent between response types indicates otherwise. Instead, this finding suggests that actions are planned prior to target presentation, and must be updated when target positions change to unexpected locations. The difference in decision time when targets are moved is on the order of 0.1 s, which is too small to account for changes in lose-shift or win-stay responding as a decay in memory of the previous reinforcement (Ivan et al, 2018).

**Table 2.** Mean decision times for lose-shift, lose-stay, win-shift, & win-stay responses. The *t*-statistic value is reported for the paired comparison between the local position changes and no change condition. Note: \**p* < .05, †*p* < .01, ‡*p* < .001 Standard errors 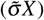 estimated from inverse-transformed RTs 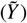 via 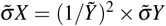.

| Local | No Global Change |  | Global Change |  |
| --- | --- | --- | --- | --- |
| | Mean±SE | *t*(105) | Mean±SE | *t*(105) |
| **Lose-Shift** |  |  |  |  |
| No Change | 0.955±.015 | - | 1.024±.017 | - |
| Displace | 0.975±.017 | 2.333* | 1.031±.018 | .936 |
| Swap | 1.025±.017 | 5.854‡ | 1.042±.019 | 1.897 |
| **Lose-Stay** |  |  |  |  |
| No Change | .980±.019 | - | 1.070±.020 | - |
| Displace | .995±.019 | 1.171 | 1.047±.018 | -1.691 |
| Swap | .987±.017 | .576 | 1.042±.019 | -2.372* |
| **Win-Shift** |  |  |  |  |
| No Change | 1.023±.020 | - | 1.084±.021 | - |
| Displace | 1.044±.021 | 1.907 | 1.081±.020 | -.287 |
| Swap | 1.065±.021 | 3.227† | 1.098±.021 | 1.447 |
| **Win-Stay** |  |  |  |  |
| No Change | 1.024±.018 | - | 1.056±.020 | - |
| Displace | 1.027±.020 | .329 | 1.064±.020 | .778 |
| Swap | 1.064±.023 | 3.644‡ | 1.058±.021 | .262 |

**Table 3.**
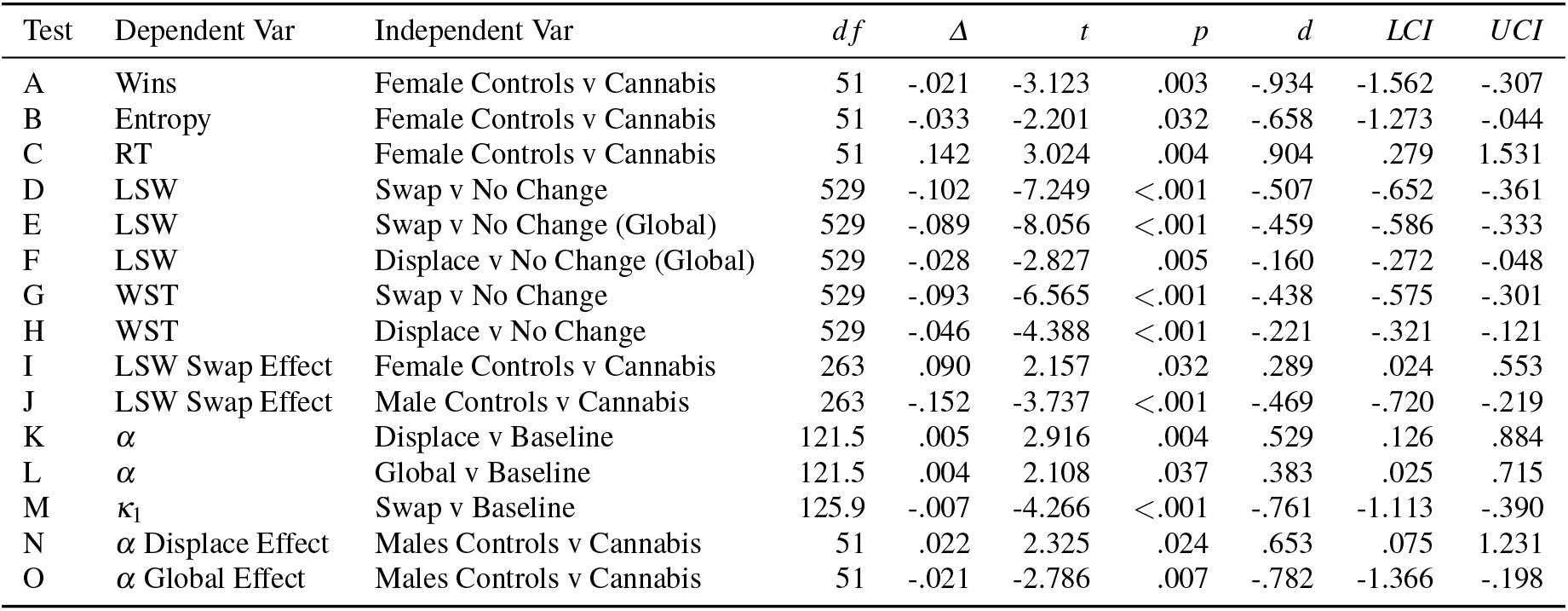
Table summarizing key significant effects. Note: *Δ* is difference between means, *d* is Cohen’s *d*, and UCI & LCI indicate boundaries of 95% confidence intervals.

### Cannabis use modulates the lose-shift

We next analyzed the effects of cannabis use on lose-shift by a mixed-effects model testing for the effects of sex (male, female), local changes (no change, displace, swap), and cannabis use (controls, habitual users). Models were fit separately to trials with and without global changes in position, in order to simplify model interpretation. A random-intercepts-only structure was used because the full random-effects structure resulted in an over-fit model.

Following no global change, there was again a significant main effect of local changes [*F*_2,1476_ = 43.347, *p* < .001] on lose-shift behaviour. Significant local × sex [*F*_2,1476_ = 5.711, *p* = .003], sex × cannabis [*F*_1,102_ = 8.342, *p* = .005], and local × sex × cannabis [*F*_2,1476_ = 13.008, *p* < .001] interactions were also present. No other effects or interactions were present (*p* > .148 in each case). As shown in Figure 4, male and female controls exhibit similar effects of positional changes on lose-shift behaviour. Both exhibit a strong-lose shift tendency that is extinguished following local swaps and with experience on the task. In females, the difference between the no change and swap conditions increases with heavy cannabis use [*t*(263) = 2.157, *p* = .032, *d* = .289]^*l*^. This effect on lose-shift behaviour may be due to an increased reliance on spatial choice cues or an increase in baseline lose-shift behaviour. We find that while female cannabis users lose-shift more in the no change condition [*t*(263) = 2.402, *p* = .017, *d* = .321], the effect of local swaps was the same [*t*(263) = -.951, *p* = .343, *d* = -.127]. In other words, female cannabis users are no more reliant on spatial choice cues than are female controls. Instead, they exhibit an elevated lose-shift response at baseline (Fig. 3.C). Conversely, men exhibited the opposite trajectory. While male controls show a large effect of spatial swaps, this behaviour is extinguished, and in some cases reversed with elevated drug use [*t*(263) = 3.737, *p* < .001, *d* = .469]^*J*^. Importantly, male cannabis users exhibited decreased lose-shifting in the no change condition [*t*(263) = −3.489, *p* < .001, *d* = -.438] and an increase following local swaps [*t*(263) = 2.494, *p* = .013, *d* = .313] relative to controls. Therefore, male cannabis users exhibit less reliance on spatial cues when responding after losses.

**Fig. 4.**
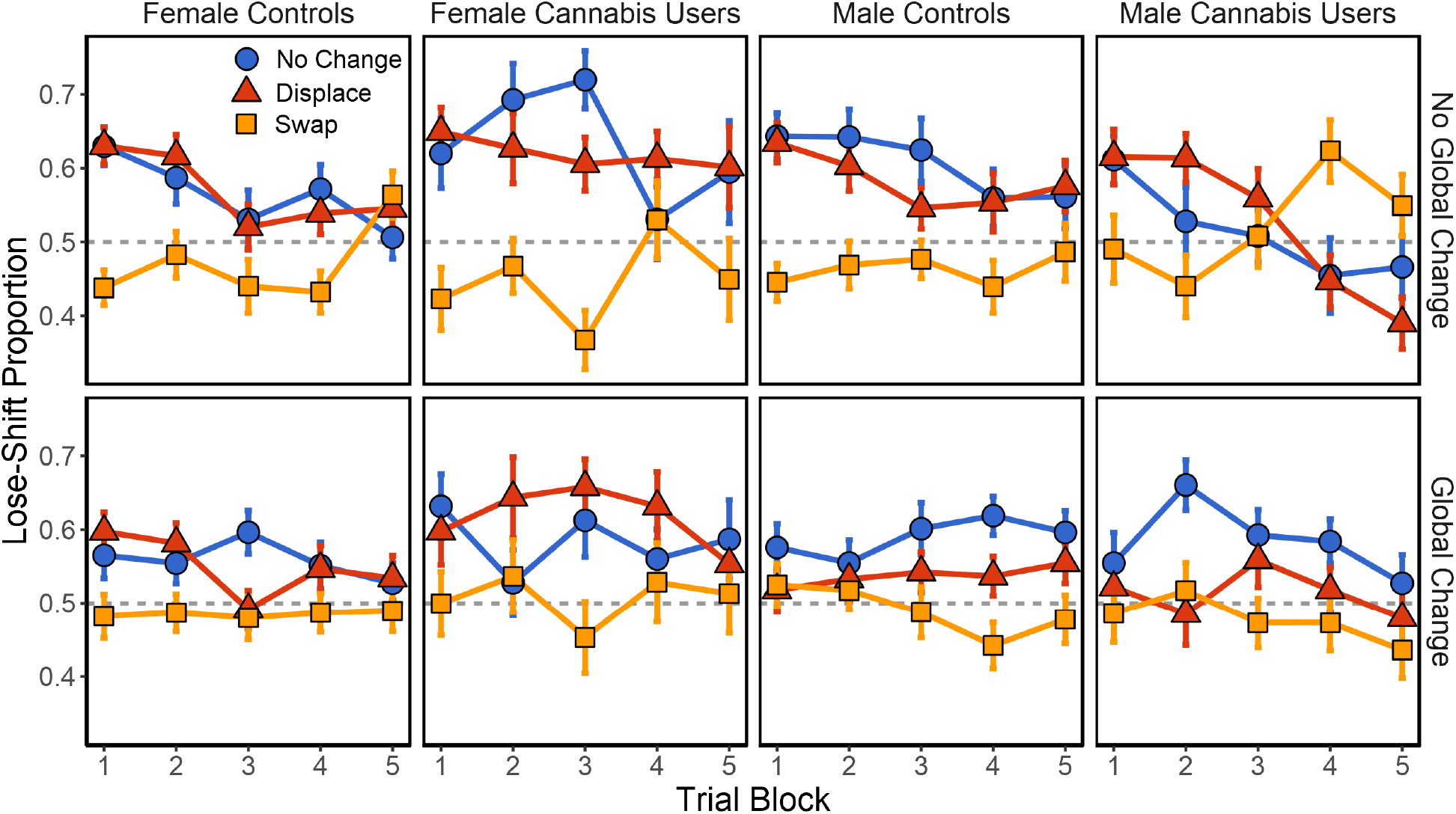
Effect of local & global changes in choice position on lose-shift tendencies in male & female cannabis users, relative to controls. SEM in error bars.

Modulation of the lose-shift response may not be specific to cannabis alone. As seen in Figure 3.E, Total drug use (ASSIST score) is positively correlated with the lose-shift swap effect in men [*r*(51) = .378, *p* = .005]. In women, however, they are uncorrelated [*r*(51) = -.143, *p* = .308], suggesting that cannabis use provides a more informative metric. Furthermore, ASSIST cannabis use score is more heavily correlated with total drug use [*r*(104) = .847, *p* < .001], than are tobacco [*r*(104) = .762, *p* < .001], or alcohol use [*r*(104) = .651, *p* < .001]. Therefore, in our population cannabis use is the most indicative of total drug use, while also remaining a clinically relevant classification.

Following global changes in position, there were significant effects of local changes [*F*_2,1476_ = 34.300, *p* < .001] on lose-shift behaviour, local × sex [*F*_2,1476_ = 5.633, *p* = .004], and sex × cannabis [*F*_1,102_ =5.764, *p* = .018], interactions. No other effects were significant (*p* > .088 in all cases).

Similar models were applied to win-stay behaviour. While the effects of local changes remained significant [*F*_2,1578_ = 22.755, *p* < .001], and those of sex were near significance [*F*_1,1578_ = 3.477, *p* = .062], the win-stay was not affected by drug use, sex, or interactions with local changes in position (*p* > .219 in all cases). As seen in Figure 3.D, processing of the win-stay did not differ with sex or drug use, and was not considered further. The same was true following global changes in position, where no effects were significant (*p* > .135 in all cases).

### Computational Results

The results presented above demonstrate that the location of choice targets, rather than their visual identity, is more important for choice adaptation based on reinforcement in the immediately previous trial. The importance of spatial configuration is evidenced by changes in win-stay and lose-shift probabilities following manipulations of cue position. We next sought to determine how choices, cue configurations, and reinforcement affected choice over multiple trials. We therefore used a biologically-relevant computational model to determine how learning rate, reward valuation, and loss aversion affected choice. Each participant’s choice behaviour was analyzed with the *Q-learning with forgetting* (FQ) model, as described by Barraclough et al (2004). It uses learning rate (*α*), inverse temperature (*β*), reward strength (*κ*_1_), and punishment strength (*κ*_2_) as parameters (hidden states) to estimate action values. We compared model performance against *Q-learning* (Q) (Sutton and Barto, 2018) and *Q-learning with differential forgetting* (DFQ) (Ito and Doya, 2009). The former only includes the *α* and *β* parameters, while the latter includes a second *α*_2_ parameter, in order to describe forgetting as a different process from learning. Hidden states were estimated for each subject, using the negative log-likelihood to assess model performance:

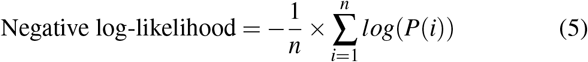

where *n* is the number of trials and *P*(*i*) the probability that the model predicted the choice made by each subject on trial *i*. As seen in Figure 5.A, both the FQ and DFQ models performed better than the Q-learning model. However, the FQ model fit no worse than DFQ (*p* = .503) while requiring one less parameter. Therefore, *Q-learning with forgetting* provided the best model of human choice in the present task. In no instance was model fit different as a result of sex or cannabis use (*p* > .154 in all cases), indicating that comparison of the parameter values is well founded. Note that the parameters were free to vary within the session, and so take a range of values for each subject. Figures 5.B-D shows a logistic-like relationship between parameter values and win-stay/lose-shift response probability. When fit against a mixed effects logistic function with random effects for logistic asymptote, intercept, and slope, we found a strong relationship between reward strength (*κ*_1_) and win-stay behaviour. Consequently, the asymptote [*β* = .370, *F*_1,30691_ = 3069.663, *p* < .001], intercept [*β* = .019, *F*_1,30691_ = 10.986, *p* = .001], and slope [*β* = -.153, *F*_1,30691_ = 237.979, *p* < .001] parameters were significant. As seen in Figure 5.B, when reward strength is low, subjects winstay at a fixed baseline of 37.0% (*SD* = 10.9%). However, when *κ*_1_ is high (> .019), subjects almost exclusively winstay. The same is true of the lose-shift. At low values of *κ*_2_ subjects exhibit a lose-stay policy. However, as *κ*_2_ increases, they reach a stable lose-shift strategy of 64.8% (SD = 10.1%). Consequently, for the lose-shift the asymptote [*β* = .648, *F*_1,32587_ = 3712.685, *p* < .001], intercept [*β* = -.033, *F*_1,32587_ = 20.921, *p* < .001], and slope [*β* = .118, *F*_1,32587_ = *p* < .001] parameters were significant. Lose-shift behaviour was also associated with learning rates (*α*, Fig. 5.D). A mixed-effects model with random asymptotes indicated the asymptote [*β* = .567, *F*_1,32587_ = 14089.972, *p* < .001], intercept [*β* = .026, *F*_1,32587_ = 3641.362, *p* < .001], and slope [*β* = .595, *F*_1,32587_ = 122.090, *p* < .001] of the relationship between *α* and lose-shifting were significant. As subjects increase the rate at which new reinforcement updates past knowledge of choice-outcome associations, they lose-shift more before reaching an asymptote of 56.7% (*SD* = 4.8%). Similar analysis of the relationship between winstay and *α* found no relationships (p > .128 in all cases).

**Fig. 5.**
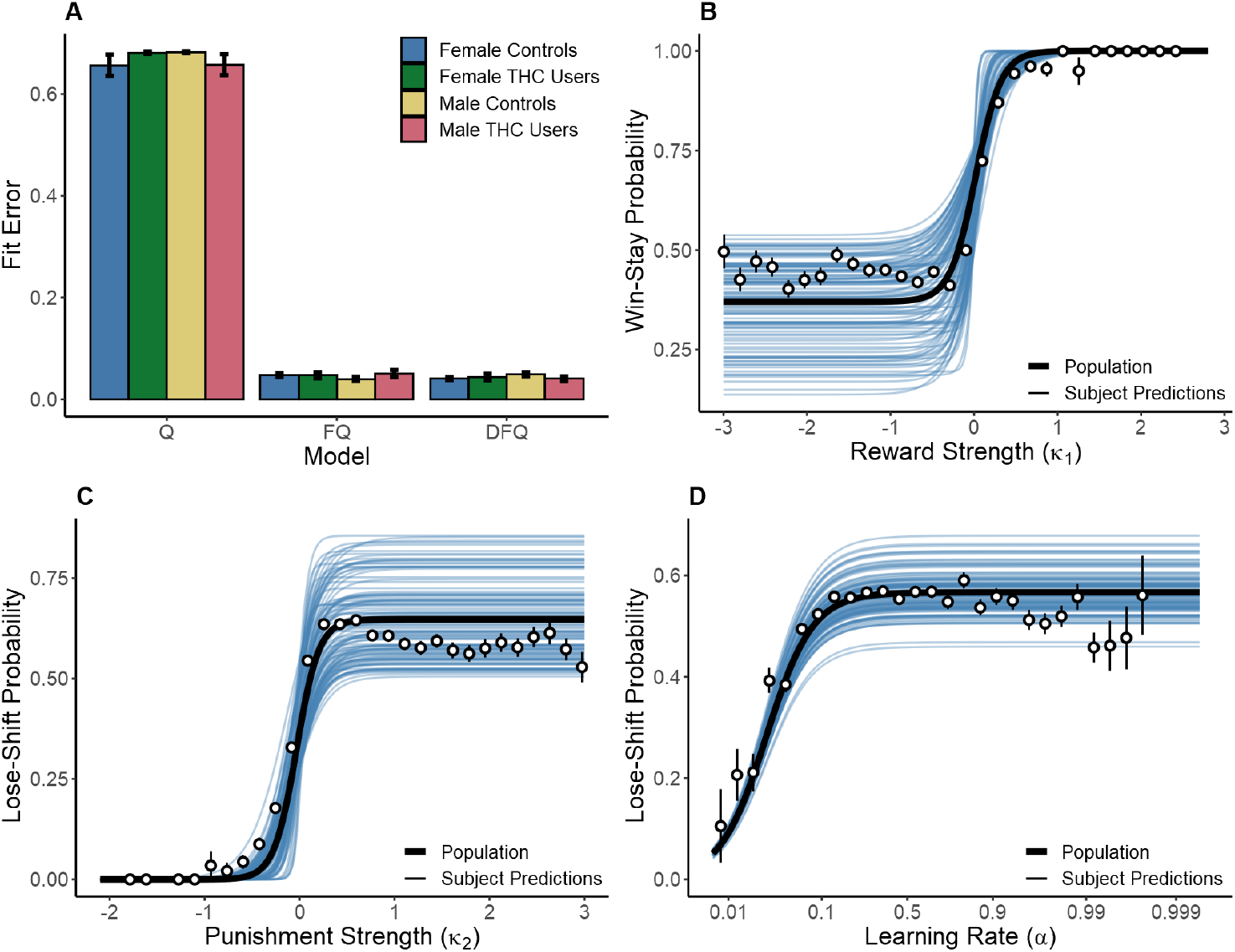
A: Performance of the *Q-learning* (Q), *Q-learning with forgetting* (FQ), and *Q-learning with differential forgetting* (DFQ) models. B: relationship between *κ*_1_ and win-stay behaviour with curves fit to individual subjects (blue lines), the population average (black line), and averages of binned raw data (points). C-D: relationship between *κ*_2_ & *α* and lose-shift behaviour. Note: *α* was normalized via the logit transform prior to model fitting.

Given the relevance of the FQ model to human behaviour, we next sought to quantify how hidden states changed in response to reinforcement and cue positions using Volterra decomposition. The method accounted for the past effects of wins, local displacement, swaps, and global changes on changes in *α*, *κ*_1_, and *κ*_2_ over the preceding *n* ∈ (1,32) trials. The effects of wins were calculated relative to those of losses, while local displacement, swaps, and global changes were calculated relative to the no change condition. Their impact on hidden states over time were tested with a mixed-effects model incorporating random intercepts and slopes for each subject. Each model was reparameterized to exclude a global intercept, but fit a separate intercept for each group (trial type). Therefore, for each model we tested whether each trial type differed from zero (null hypothesis of no effect) to determine if it had a significant impact on RL parameters.

Initially we collapsed data over sex and cannabis use to determine what variance between subjects is explained by the model. For learning rate (*α*), there was a significant effect of trial type (of past trials) on the learning rate of the present trial [*F*_4,404.68_ = 7.828, *p* < .001]. There was a significant change from baseline following local displacement [*t*(121.5) = 2.916, *p* = .004, *d* = .529]^*K*^ or global changes [*t*(121.5) = 2.108, *p* = .037, *d* = .383]^*L*^, but not local swaps or winning outcomes (*p* > 0.091 in both cases). As seen in Figure 6.A, all trial types resulted in a slight increase in learning rates relative to baseline (Volterra weight intercept > 0). The plot indicates that the effect persists for a maximum of about 10 previous trials.

**Fig. 6.**
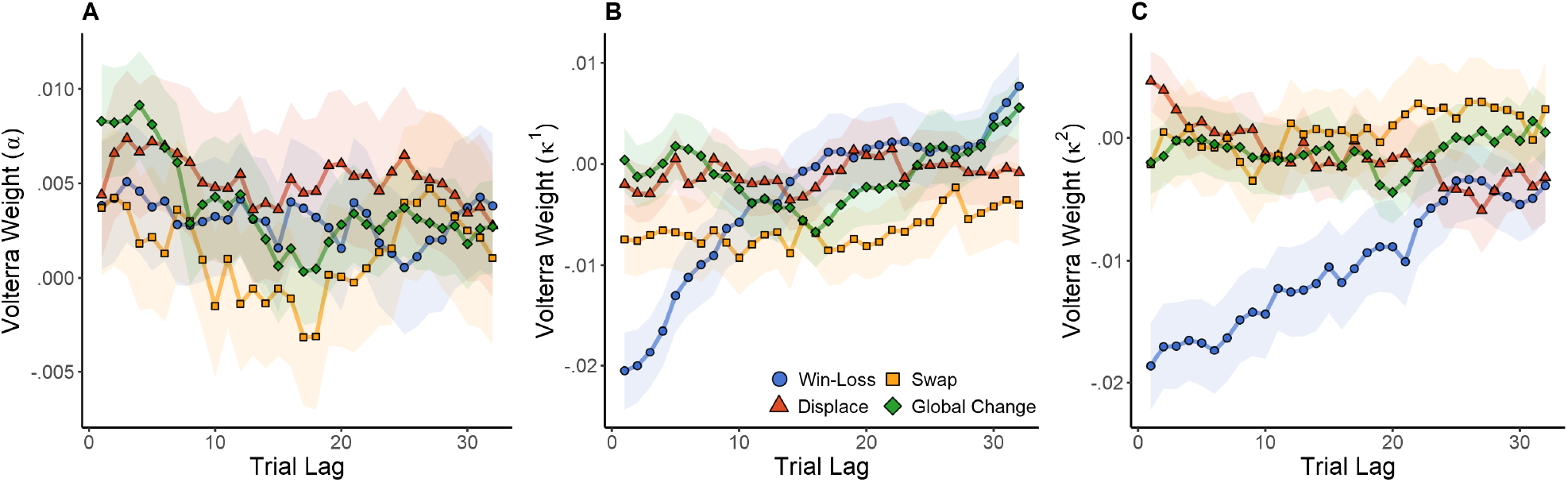
Effect of wins, local, & global changes in choice position on Q-learning parameters *α* (learning rate), *κ*_1_ (reward strength), & *κ*_2_ (punishment strength). The influence of wins and cue rearrangement during the previous 32 trials is estimated by Volterra decomposition, which provides a weight (loading) for each trial lag. SEM in shaded area.

There was also a significant effect of past trial type on reward strength [*F*_4,404.68_ = 19.629, *p* < .001]. Wins [*t*(125.9) = −2.209, *p* = .029, *d* = -.394] and local swaps [*t*(125.9) = −4.266, *p* < .001, *d* = -.761]^*M*^ both caused significant decreases in reward strength (Volterra weight intercept < 0). Therefore, as multiple wins (or swaps) are experienced, future rewards become progressively less impactful on choice. Local displacement and global changes had no effect on reward strength (*p* > .461 in both cases). Punishment strength (*κ*_2_) was also affected by trial type [*F*_4,404.68_ = 67.857, *p* < .001]. As with *κ*_1_, wins (relative to losses) decreased the strength of future punishments [*t*(142.4) = −8.025, *p* = .029, *d* = −1.345]. Consequently, losses increased the strength of future punishment, so that experiencing multiple losses would have a cumulative effect. As seen in Figure 6.B & C, reward strength quickly recovered in response to wins. However, *κ*_2_ exhibited a much more prolonged change, suggesting that the effects of losses were more impactful over a longer time course. No other trial type had a significant effect on *κ*_2_ (*p* > .321 in all cases).

In sum, these data indicate that recent rewards and manipulation of choice target locations increase the learning rate. Wins reduce the sensitivity of subjects to future reward (*κ*_1_) and punishment (*κ*_2_), whereas losses increase the sensitivity. We interpret this to indicate that subjects who have been winning on recent trials persist in their long-term strategy (e.g. executive control), rather than engaging in reflexive responding strongly influenced by the immediately previous reinforcement (e.g., sensorimotor control).

We next tested whether cannabis use and sex modulated the response of reinforcement learning parameters to wins, local displacements, swaps, and global changes. We used a mixed effects model with random slopes and intercepts for each subject. In this case, a global intercept was used because we were testing differences between conditions, rather than between each group relative to the null hypothesis of no change within each condition.

Cannabis use and sex had a significant effect on the change in learning rates (*α*) following local displacement, as evidenced by a cannabis × sex interaction [*F*_1,102_ = 4.748, *p* = .032], while there were no main effects of sex or cannabis use (*p* > .178). The same sex × cannabis interaction was also present in the response to global changes [*F*_1,102_ = 7.443, *p* = .007]. However, there were no differences in the response to winning outcomes or local swaps (*p* > .108 in all cases). The immediate responses to each trial type (in the following trial, or at lag=1) are shown in Figure 7.A. Males exhibit a significant increase in learning rates immediately following local displacement [*t*(51) = 2.325, *p* = .024, *d* = .653]^*N*^. Therefore, local displacement increases the rate at which new information updates choice value estimates. Male cannabis users exhibited a similar increase in response to local swaps, though this fell short of significance [*t*(51) =1.801, *p* = .078, *d* = .506]. Conversely, learning rates fell in response to global changes for male cannabis users, relative to male controls [*t*(51) = −2.786, *p* = .007, *d* = -.782]^*O*^. For *κ*_1_, there was a significant effect of sex on the response to displacement [*F*_1,102_ = 4.517, *p* = .036], as seen in Figure 7.B. In addition, there was a significant sex × cannabis interaction in the effect of global changes on *κ*_1_ [*F*_1,102_ = 6.242, *p* = .014]. However, for *κ*_2_, male and female cannabis users did not differ from controls in their response to wins, local displacement, swaps, and global changes (*p* > .138 in all cases).

**Fig. 7.**
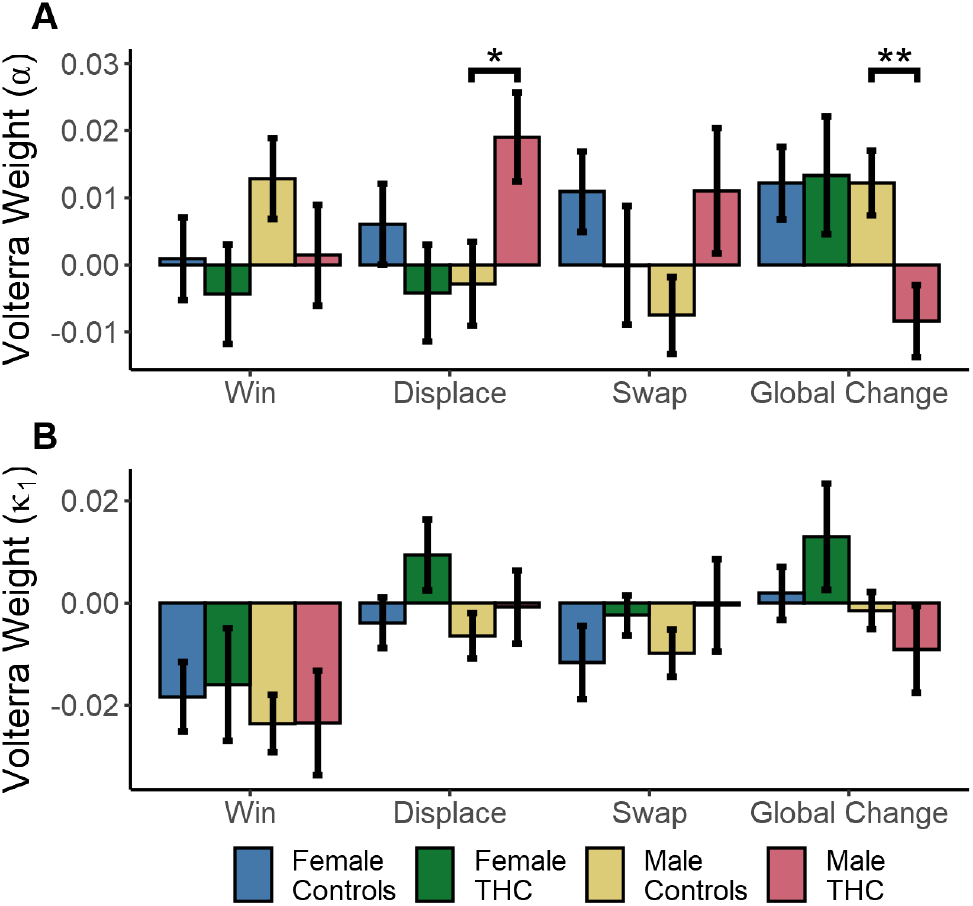
Cannabis × Sex interactions on mean reinforcement learning parameter values estimated by Volterra decomposition. Learning rate *α* (A) & effect of wins *κ*_1_ (B) to wins, local displacement, swaps, and global changes. SEM in error bars.

In sum, male cannabis users tended to increase learning following local, but not global changes of target positions, which is different from all other groups. Moreover, the parameter values for female cannabis users were not different from controls, suggest that their reduced task performance is related mostly to the processing of the previous reinforcement (e.g. lose-shift) rather than effects spanning multiple trials.

## Discussion and Conclusions

The data here provide novel behavioural evidence that the lose-shift response is a product of sensorimotor systems, and that the regulation of such responding is compromised differently in men and women with high recreational cannabis use. A high proportion of lose-shift responding is sub-optimal in the present task because it is predictable, and can therefore be exploited by the computer opponent. Indeed, the propensity for lose-shift responding is negatively correlated with task performance here. Nonetheless, subjects engage this response above chance levels for several hundred trials before learning to suppress it. This suggests that it is a default strategy, consistent with previous work in humans (Ivan et al, 2018), and analogous to what has been observed in animals performing a similar task (Gruber and Thapa, 2016). As lose-shift responses eventually converge to chance levels in trials with no global changes, the probability of win-stay responses increase above chance levels. We found that this negative correlation between lose-shift and winstay was significant and strikingly similar to rodents (Gruber and Thapa, 2016), suggesting that lose-shift and winstay are expressed by neural systems in competition with one another.

We show here that participants overwhelmingly perform the lose-shift according to target position, rather than target identity. In other words, participants avoided the prior position of the previously chosen target when it was unrewarded. This novel observation reveals a strong spatial component to the lose-shift. These data are consistent with the notion that lose-shift is a product of sensorimotor systems. Loss-driven response shifting is reduced following lesions to sensorimotor striatum in animals (Skelin et al, 2014; Gruber et al, 2017; Thapa and Gruber, 2018) or damage to putamen/insula in humans (Danckert et al, 2011); these homologous structures are strongly involved with sensorimotor control. Moreover, decision times are lower for lose-shifting than for lose-stay responses, even when global changes in position (that require equally distant arm movements) are present. There are multiple reasons this may occur. First, the visual systems of the brain process information about spatial position independently from other object characteristics (Mishkin et al, 1983; Haxby et al, 1991). The ventral ‘where’ pathway processes information more quickly than the ‘what’ pathway. (Deubel et al, 1998; Goodale and Milner, 1992). Secondly, the ventral pathway may be used to compute actions prior to stimulus presentation. In the perceptual learning literature, activity in both the motor and visual cortices builds up prior to stimulus onset, and reflects stimulus expectation and the associated motor responses (de Lange et al, 2013). Moreover, pre-response fluctuations in beta-power motor activity are also predictive of choice alternation (i.e., lose-shift), regardless of associated motor action (Pape and Siegel, 2016). There is evidence that loops involving premotor cortex and the lateral striatum map vision and other sensory modalities into an egocentric space. The ventral premotor cortex contains neurons that both drive motor actions, but also encode locations of visual, tactile, and auditory stimuli (Fadiga et al, 2000). Consequently, they form a motor vocabulary for mapping several modalities into an actions in a common egocentric space. Even when stimuli are removed, these neurons still respond to the position of remembered objects in relation to the body (Graziano and Gross, 1998). The putamen (LS in rodents) also contains these bimodal visuomotor cells (Graziano and Gross, 1996), and therefore has the capacity to mediate lose-shift from a remembered location. In the context of our study, spatial rearrangement of choice targets subverts this motor preparation, requiring choices to be recalculated following stimulus onset. This is evidenced by the increase in response times following local swaps and displacement. Interestingly, local swaps had a larger and more consistent effect on response times than did displacement, suggesting that a greater level of motor recalculation is required. Specifically, we speculate that it requires more time for the executive control to overcome the intrinsic inhibition of a previously unrewarded action (than a novel one) in the motor system in order to intentionally select it.

The influence of local and global changes in choice position also highlights the importance of egocentric and allocentric processing of space. While local changes in choice orientation modulate the lose-shift, these effects persist even when all choices are moved to a new global position relative to the observer. Conversely, the win-stay is much less affected by spatial position. Local changes do have an effect on behaviour, but these are eliminated by concurrent global changes. These results highlight the importance of allocentric processing on the lose-shift. Choice positions are calculated relative to one another, allowing their associated values to be maintained across large global movements in choice position. Conversely, while processing of the winstay is less reliant on spatial information, egocentric reference frames are more important than allocentric, where choice value is calculated relative to the subject. Consequently, local and global changes have a large effect on win-stay behaviour.

In addition to driving different decision strategies, the putamen/LS and ventral striatum (VS, including the nucleus accumbens) also respond differently to psychoactive drugs. Relative to the VS, the LS exhibits a much higher density of dopamine transporters (Coulter et al, 1997), endocannabinoid receptors (Herkenham et al, 1991), opioid receptors (Benfenati et al, 1991), and alcohol-sensitive NMDA receptors (Liste et al, 1995). Consequently, THC administration temporarily increases dopamine release throughout the striatum, but particularly in the LS (Jentsch et al, 1998; Sakurai-Yamashita et al, 1989). The effects of acute ethanol exposure are similar, though greater in the VS (Vena et al, 2016; Clarke et al, 2015). Conversely, long-term sensitization to alcohol and cannabis reduces availability of striatal dopamine receptors (Budygin et al, 2007; Martinez et al, 2005; Albrecht et al, 2013) and cannabinoid receptors (Villares, 2007), especially in the LS. Chronic exposure also inhibits the prefrontal cortex, shifting choice control to the LS (Lucantonio et al, 2014; Everitt and Robbins, 2005, 2013). We expect this effect to impair the ability of participants to use executive control to suppress lose-shift responding by the sensorimotor systems. We do not have sufficient primary evidence to hypothesize how the change in receptor densities by repeated alcohol/THC exposure affects lose-shift processing within the LS and/or other components of the sensorimotor system.

In the present study, we find that self-reported level of recreational use of cannabis affects task performance, but that this differs on the basis of biological sex. Elevated cannabis use in men decreased spatial modulation of the lose-shift, possibly through dopaminergic desensitization of the LS. As seen in Figure 4, baseline lose-shifting is also reduced, falling below 50% in trial blocks 4 and 5. With this reduction, lose-shift responding after swaps increases to 63%. Therefore, either the calculation of the lose-shift is affected or its suppression by executive systems is enhanced in male cannabis users, while spatial processing remains unaffected. Conversely, female cannabis users exhibit decreased task performance and choice entropy - behaviours thought to rely on the suppression of sensorimotor responding by the prefrontal cortex. Furthermore, they show a moderate and significant increase in baseline lose-shift responding [*F*_1,51_ = 4.109, *p* < .048], revealed by a mixed effects model between female controls and cannabis users in the no change condition (3.C). It thus appears that females with high cannabis use exert less executive control over sensorimotor systems in our task.

While it is tempting to describe this sex difference as a consequence of different drug effects on the brain in males and females, we cannot strongly support this inference based on the present data. Several alternatives are possible. It is possible that the effect is due to a confounding factor that promotes high levels of recreational drug use and also impairs sensorimotor regulation. Unfortunately, the WHO ASSIST is not sufficient to infer whether these are the case in the present study. However, it is known that females are more susceptible to drug tolerance (including cannabis) and sensitization than are males (Wakley et al, 2014; Robinson, 1988). Drug use is also comorbid with mood and anxiety disorders, particularly depression (Zilberman et al, 2003), which causes heightened loss aversion (Beevers et al, 2013). These differences are possibly due to the effects of estrogen, which enhances striatal dopamine release in response to psychoactive drugs (Becker, 1999) and alters the effects of drugs on the prefrontal cortex. Females rats with high estrogen levels exhibit dysfunction of the prefrontal cortex relative to males and low-estrogen females when exposed to dopamine-enhancing drugs (Shansky et al, 2004). Estrogen also heightens the effects of cocaine and amphetamine, causing an abnormal BOLD response in rats (Sárvári et al, 2014; Febo et al, 2005). Alcohol and cannabis consumption also increase oestradiol levels, and can inhibit testosterone production in males (Yonker et al, 2005; Purohit, 2000; Maskarinec et al, 1978; Kolodny et al, 1974; Harclerode, 1984). In males, increased estrogen and reduced testosterone levels cause declines in spatial cognitive ability Janowsky et al (1994). Therefore, the heightened susceptibility of the PFC to the combined effects of estrogen and drug abuse provides an explanation for why only women with high ASSIST scores show a dominance of sensorimotor control, without compromising the spatial dependence of lose-shift. Specifically, this population had accelerated decision speeds, lower proportion of wins, and a tendency for lower entropy of response sequences. On the other hand, the lose-shift remained sensitive to swapping cue locations, which is similar to controls, but opposite of what is observed in males with high ASSIST scores. Our analysis of behaviour through a reinforcement learning framework also revealed a cannabis × sex interaction. Whereas the other analysis presented here focuses on the effects of the previous trial, the Q-learning model allowed us to examine effects that span many trials. It was the men who used cannabis who stood out in this analysis; they had increased learning when previous cues were displaced locally, and de-creased learning when previous cues were switched globally. We expect such learning is part of a reinforcement learning scheme in ‘goal-directed’ brain circuits linked more closely to executive function than sensorimotor control (Balleine and O‘Doherty, 2010; Gruber and McDonald, 2012), suggesting that not only is there an enhanced suppression of sensorimotor control by executive function in male cannabis users, but that adaptation by the executive system is also different than the other groups. It is worth noting, however, that our sample (as is common in the field) was predominantly young university students, who are presumably well educated and high functioning. We urge caution in extrapolating our findings to the general public.

The interpretation of data in this study faces several challenges besides the aforementioned limitations of the ASSIST. First, alcohol and cannabis use are highly concordant (Spearman’s correlation of *ρ*=.533, *p* < .001 in our sample), and likely additive in their effects. Second, the sexually dimorphic effects observed here may be due to confounding interactions between drug use, IQ, and/or psychiatric disorders that have different prevalence among the sexes. However, the sexually dimorphic distribution of endocannabinoid receptors in the striatum and prefrontal cortex (De Fonseca et al, 1994) likely also play an important role. For instance, errors when reconstructing spatio-temporal sequences were reduced in men and increased in women following THC treatment (Makela et al, 2006). We previously reported that lose-shift is decreased by acute administration of THC in female rats (Wong et al, 2017a). It is possible that the down regulation of receptors in have users may cause the inverse, which would be consistent with the data here.

In sum, the data presented here indicate that lose-shift responding is a useful gauge of the cognitive control over sensorimotor responding in humans, and that this is impacted differently in men and women that heavily use cannabis. These linkages are important factors to account for the impact of lose-shift responding in real-world economic decision making, such as gambling (Abouzari et al, 2015; Worthy et al, 2013), as well as clinical/laboratory testing of cognitive flexibility with tasks such as the Wisconsin Card Sorting Task that involve loss-based shifting of response policies.

## Acknowledgements

We would like to thank Donna Waxman for helping with data collection.

